# Automation of (Macro)molecular Properties Using a Bootstrapping Swarm Artificial Neural Network Method: Databases for Machine Learning

**DOI:** 10.1101/779496

**Authors:** Blerta Rahmani, Hiqmet Kamberaj

## Abstract

In this study, we employed a novel method for prediction of (macro)molecular properties using a swarm artificial neural network method as a machine learning approach. In this method, a (macro)molecular structure is represented by a so-called *description vector*, which then is the input in a so-called *bootstrapping swarm artificial neural network* (BSANN) for training the neural network. In this study, we aim to develop an efficient approach for performing the training of an artificial neural network using either experimental or quantum mechanics data. In particular, we aim to create different user-friendly online accessible databases of well-selected experimental (or quantum mechanics) results that can be used as proof of the concepts. Furthermore, with the optimized artificial neural network using the training data served as input for BSANN, we can predict properties and their statistical errors of new molecules using the plugins provided from that web-service. There are four databases accessible using the web-based service. That includes a database of 642 small organic molecules with known experimental hydration free energies, the database of 1475 experimental pKa values of ionizable groups in 192 proteins, the database of 2693 mutants in 14 proteins with given values of experimental values of changes in the Gibbs free energy, and a database of 7101 quantum mechanics heat of formation calculations.

All the data are prepared and optimized in advance using the AMBER force field in CHARMM macromolecular computer simulation program. The BSANN is code for performing the optimization and prediction written in Python computer programming language. The descriptor vectors of the small molecules are based on the Coulomb matrix and sum over bonds properties, and for the macromolecular systems, they take into account the chemical-physical fingerprints of the region in the vicinity of each amino acid.

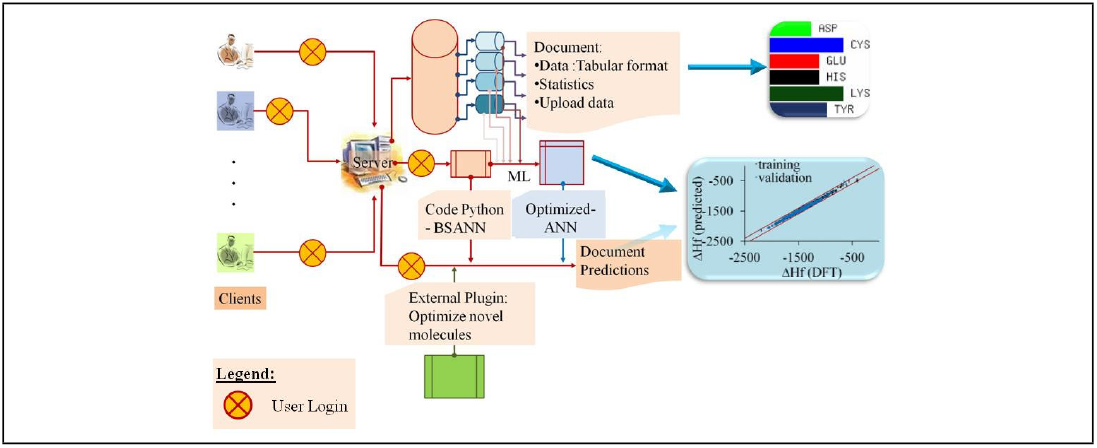
Graphical TOC Entry.

## 1 Introduction

Recently, neural network method has seen a broad range of applications in molecular modeling. ^1^ In Ref.,^2^ a hierarchical interacting particle neural network approach is introduced using quantum models to predict molecular properties. In this approach, different hierarchical regularization terms have been introduced to improve the convergence of the optimized parameters. While in Ref.,^3^ the machine learning like-potentials are used to predict molecular properties, such as enthalpies or potential energies. The degree to which the general features included in characterizing the chemical space of molecules to improve the predictions of these models is also discussed in Refs.^4,5^ Tuckerman and co-workers^6^ used a stochastic neural network technique to fit high-dimensional free energy surfaces characterized by reduced subspace of collective coordinates. While very recently^7^ a comparison study has been performed between neural network approach and Gaussian process regression to fit the potential energy surfaces. One of the recognized problems in using machine learning approaches in prediction free energy surfaces is the inaccurate representation of general features of the surface topology by the training data. To improve on this, a combination of metadynamics molecular dynamics with neural network chemical models have also proposed.^8^ It is worth noting that in the prediction of free energy surfaces, an accurate representation of the reduced subspace can be important. For that, Wehmeyer & Noé^9^ have used the time-lagged auto-encoder to determine essential degrees of freedom of dynamical data.

Machine learning approaches have also been in the field of drug-design, for instance, in predicting drug-target interactions,^10^ and it is a promising approach. In particular, the method is used in combination with molecular dynamics to predict the ligand-binding mechanism to purine nucleoside phosphorylase,^11^ and it accurately identifies the mechanism of drug-target binding modes.

In this study, we employed a novel method for prediction of (macro)molecular properties using a swarm artificial neural network method as a machine learning approach. In this method, a (macro)molecular structure is represented by a so-called *description vector*, which then is used as input in a so-called *bootstrapping swarm artificial neural network* (BSANN) for training the neural network. We aim to develop an efficient approach for performing the training of an artificial neural network using either experimental or quantum mechanics data. In particular, we created different user-friendly online accessible databases of well-selected experimental (or quantum mechanics) results that can be used as proof of the concepts. Furthermore, with the optimized artificial neural network using the training data served as input for BSANN, we can predict properties and their statistical errors of new molecules using the plugins provided from that web-service.

There are four databases accessible using the web-based service. The database of 642 small organic molecules with known experimental hydration free energies, ^12^ which well-studied in Ref.;^13^ the database of 1475 experimental pKa values of ionizable groups in 192 proteins (including 153 wild-type proteins and 39 mutant proteins);^14–17^ the database of 2693 mutants in 14 proteins with given values of experimental values of changes in the Gibbs free energy;^18,19^ and a database of 7101 quantum mechanics heat of formation calculations with the Perdew-Burke-Ernzerhof hybrid functional (PBE0).^5,20^

All the data are prepared and optimized in advance using the AMBER force field^21^ in CHARMM macromolecular computer simulation program.^22^ The BSANN is the code for performing the optimization and prediction written in Python computer programming language. The descriptor vectors of the small molecules are based on the Coulomb matrix and the sum over bonds properties, and for the macromolecular systems they take into account the chemical-physical fingerprints of the region in the vicinity of each amino acid.

## 2 Materials and Methods

### 2.1 Artificial Neural Network

Machine Learning (ML) approach provides a potential method to predict the properties of a system using decision-making algorithms, based on some predefined features characterizing these properties of the system. There exist different ML methods used to predict missing data or discover new patterns during the data mining process.^23^ Neural networks method considers a large training dataset, and then it tries to construct a system, which is made up of rules for recognizing the patterns within the training data set by a learning process.

In general, for an ANN with *K* hidden layers (see also Figure 1), the output *Y_i_* is defined
as

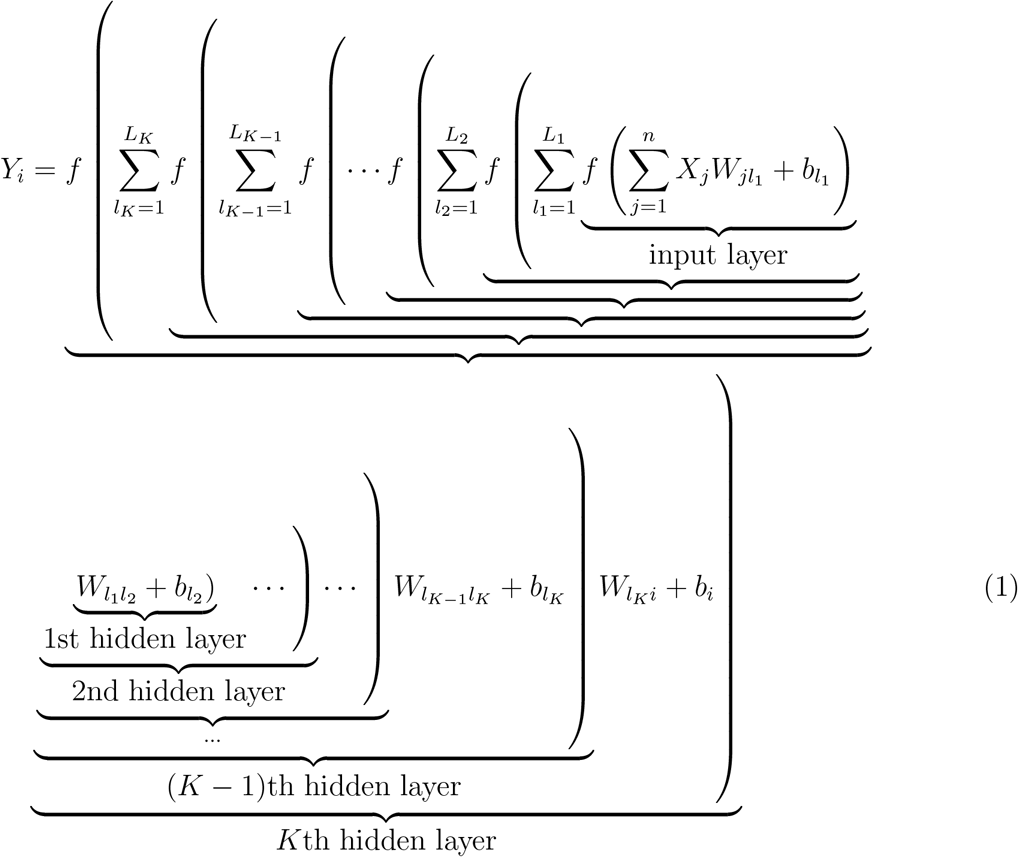

**Figure 1:**
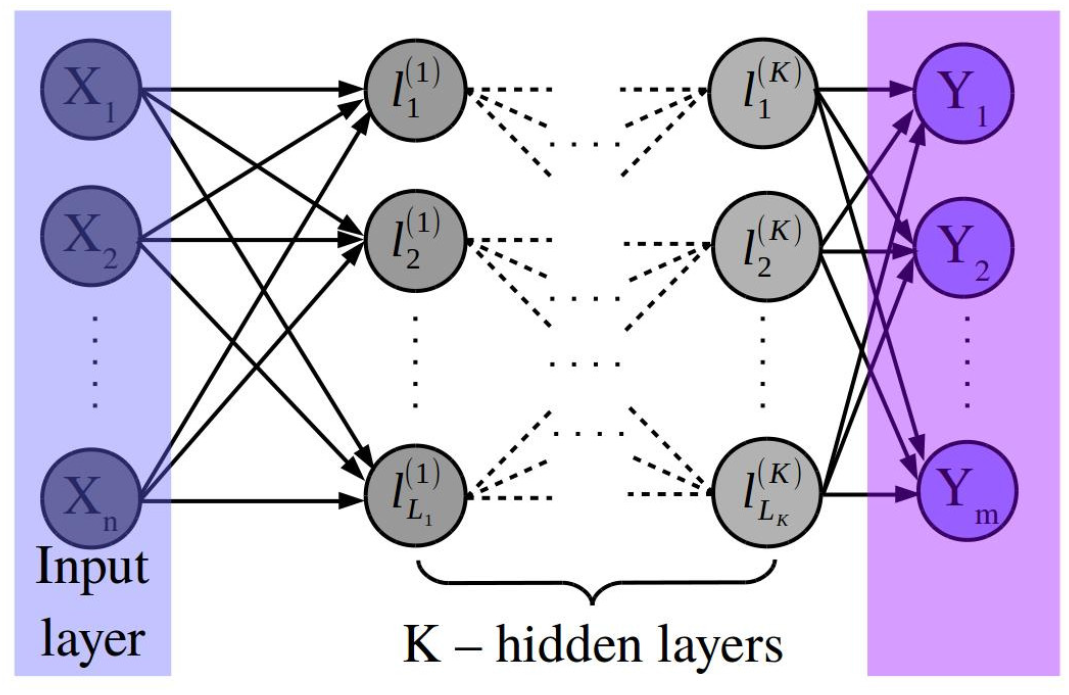
Illustration diagram of an artificial neural network (ANN). It is characterized by an input vector of dimension *n, K* hidden layers of 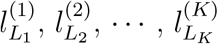 neurons each, and an output vector of dimension *m*.^24^

Here, **W** and **b** are considered as free parameters, which need to be optimized for a given training data used as inputs and given outputs, which are known. To optimize these parameters the so-called loss function is minimized using Gradient Descent method:^25^

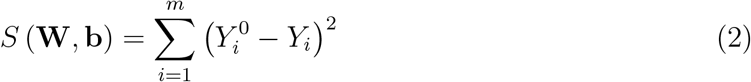

where **Y**^0^ represent the true output vector. For that, the gradients of *S* (**W**, **b**) with respect to **W** and **b** are calculated: ^25^

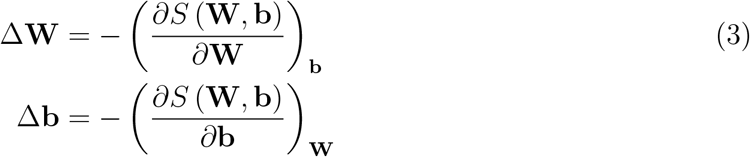

To avoid over-fitting, which is one of pitfalls of the machine learning approaches,^26^ the following regularization terms have been introduced in literature:

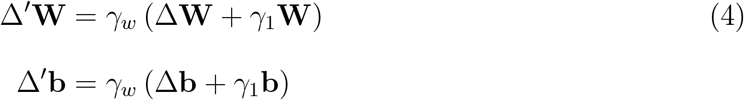

where *γ_w_* is called learning rate for the gradient and *γ*_1_ is called the regulation strength.

Usually, the Gradient Descent method often converges to a local minimum, and hence it provides a local optimization to the problem. To avoid that a new Bootstrapping Swarm Artificial Neural Network method is proposed in the literature, ^24^ which is introduced in the following.

### 2.2 Bootstrapping Swarm Artificial Neural Network

The standard ANN method deals with random numbers, which are used to initialize the parameters **W** and **b**; therefore, the optimal solution of the problem will be different for different runs. In particular, we can say that there exists an uncertainty in the calculation of the optimal solution (i.e., in determining **W** and **b**.) To calculate these uncertainties in the calculation of the optimal parameters, **W** and **b**, we introduce a new approach, namely bootstrapping artificial neural network based on the method proposed by Gerhard Paass,^27^ or similar methods.^28^ In this approach, *M* copies of the same neural network are run independently using different input vectors. Here, we implement that at regular intervals, to swap optimal parameters (i.e., **W** and **b**) between the two neighboring neural networks, which is equivalent to increasing the dimensionality of the problem by one; that is, if the dimensionality in each of the replicas is *d* = *K* × *L*, then the dimensionality of the bootstrapping artificial neural network based on the method is *d* + 1. Figure 2 shows the layout of this configuration.

**Figure 2:**
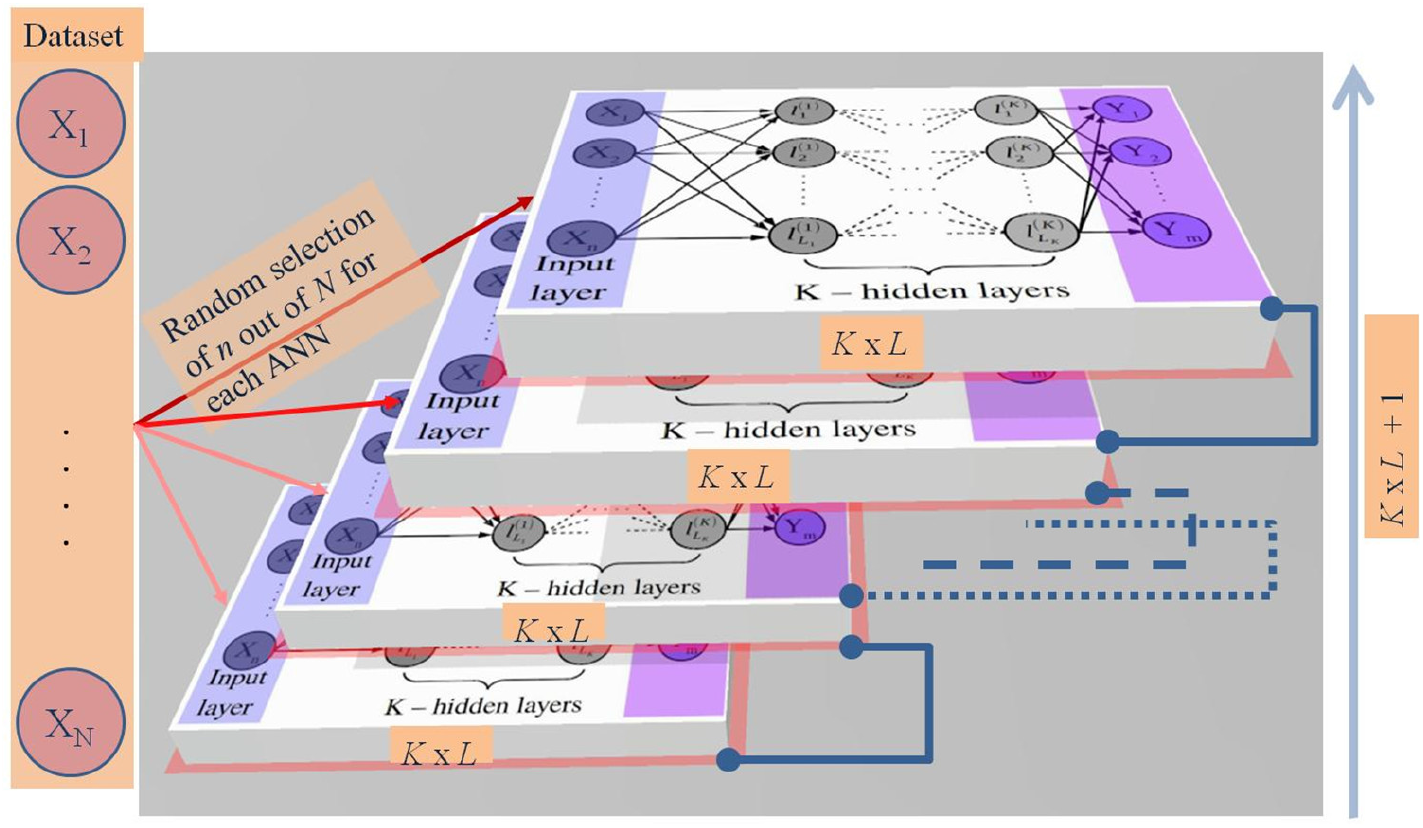
Layout of the Bootstrapping Swarm Artificial Neural Network (BSANN) as adopted by.^24^ It is characterized by *M* different input vectors each of dimension *n, K* hidden layers of 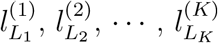 neurons each, and *M* different output vectors each of dimension *m*. Every two neighboring neural networks communicate regularly with each other by swapping the optimized parameters.

Furthermore, to achieve a good sampling of the phase space extended by the vectors **W** and **b**, we introduce two other regularization terms similar to the swarm-particle sampling approach. First, we define two vectors for each neural network, namely 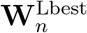 and 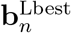, which represent the best local optimal parameters for each neural network *n*. In addition, we also define **W**^Gbest^ and **b**^Gbest^, which represent the global best optimal parameters among all neural networks.

Then, the expressions in Eq. 4 are modified by introducing these two regularization terms as the following:

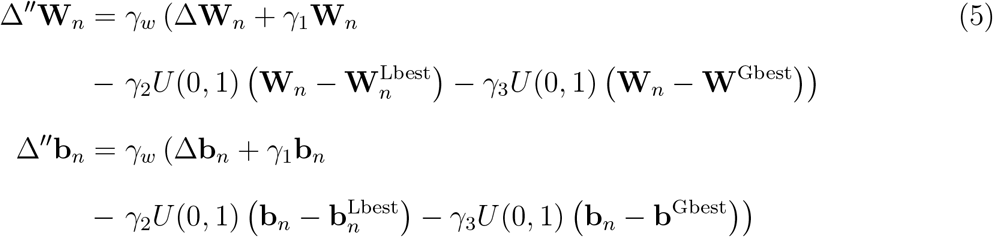

for each neural network configuration *n, n* = 1, 2, ⋯, *M*. Here, *U*(0, 1) is a random number between zero and one, and *γ*_2_ and *γ*_3_ represent the strength of biases toward the local best optimal parameters and global best optimal parameters, respectively. The first term indicates the individual knowledge of each neural network and the second bias term the social knowledge among the neural networks. This method is called here, Bootstrapping Swarm Artificial Neural Network (BSANN). Then, the weights, **W***_n_*, and biases, **b***_n_*, for each neural network *n* are updated at each iteration step according to:

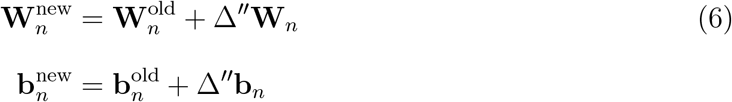

### 2.3 (Macro)molecular Feature Description

To construct data-driven models, such as in the ML approach, we will need to specify a list of physical and chemical properties of the input (macro)molecule that contain necessary information about the system. Here, the input data will be presented by a vector of length *N*, called **X**. That process is called *feature description*, and the input data are called *feature descriptors*.

Often, a so-called *Simplified Molecular Input Line Entry System* (SMILES) is used to represent a small molecule as a string of letters.^29^ In such case, the atoms could be encoded by a single integer number, such as H= 1, C= 2, N= 3, and so on, or by the nuclear charge *Z*, such as H= 1, C= 6, N= 7, and so on.^30^ It can be seen that it creates an (unnecessary) relationship between the input data, namely H<C<N, which could influence on the network performance. Other encoding models are also suggested, for instance, representing each atom of the input molecule by the following fingerprint: H= [1 0 0 ⋯], C= [0 1 0 ⋯], N= [0 0 1 ⋯], and so on.^30^ However, these fingerprints do also have drawbacks because the dimensions of the encoding vector depend on the number of atoms in the structure and may vary from molecule to molecule; also, based on this model, the atoms belonging to the same group in periodic table of elements do not behave the same.

In this study, we used the so-called *Coulomb matrix*, **C**, to encode the molecular features, which contains both the geometrical information of the three-dimensional structure and the atom type in the molecule.^20^ For any two atoms *i* and *j* in a given input molecule, the matrix element *C_ij_* is defined as:

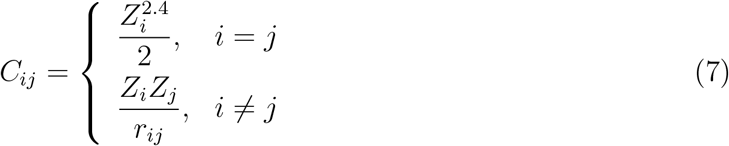

where *Z_i_* is the atomic number of the *i*th atom and *r_ij_* is the distance between the atoms *i* and *j*. The fingerprint represented by the Coulomb matrix, **C**, has some advantages, such as it takes into account the three-dimensional molecular structure, and it is invariant under rotational and translation of the structure. To calculate **C** for a given molecular structure, we need the nuclear charges for each atom and the Cartesian coordinates of the atomic positions taken from the equilibrium geometry. However, note that **C** is not invariant under the permutations of the atom order in molecule. Therefore, the spectrum of eigenvalues of matrix **C** can be used as a fingerprint of the molecule, since they are invariant under both rotation/translation and permutations of the rows and columns. A second feature descriptor that we used in this study is the so-called *sum over bonds*, which is a numerical descriptor representing the vector of bond types present in a molecule, similar to.^31^ If *N_b_* is the number of unique bonds in the dataset of the compounds studied, then a vector with dimensions *N_b_* is constructed for each molecule with entry either zeros or the integers giving the frequency of appearance for each bond type in molecular structure. This fingerprint descriptor vector has a unique length within the dataset. Then, the vector descriptor of the sum over bonds is concatenated at the end of the Coulomb matrix descriptor. To construct the input descriptor vector for a macromolecule, we introduced the following model. For example, suppose we would like to calculate the change on the Gibbs free energy upon the mutations (either single or multiple mutations) or perform pKa calculations for a selected residue in a protein. We label each residue or nucleotide of the input sequence with an ID from 1 to 24. That is, we form a descriptor vector with length *N*_1_ = 24, **X**_1_, which is a vector of zeros and ones defined as the following:

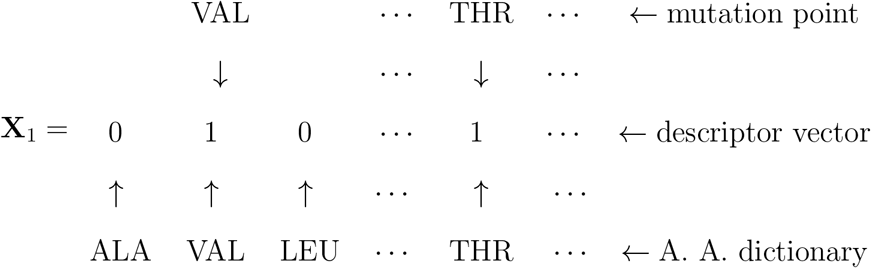

where *A. A. dictionary* represents the dictionary of the all amino acids. In addition, to characterize the environment around any mutation point, we determine another descriptor vector, namely **X**_2_ with length *N*_2_ = 24, which is defined as the following. For each mutation point amino acid i, we determine the nearest neighbor amino acids *k* = *i*_1_, *i*_2_, ⋯, *i_n.n._*, based, for example, on the center of mass distance. Then, the *j*th element 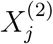 of the vector **X**_2_ is defined as a modified ‘Coulombic matrix’:

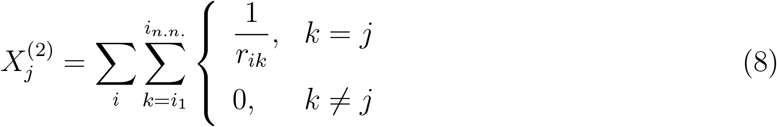

where the first sum runs over all point mutation amino acids, and the second sum runs over all nearest neighbors of amino acids *i*. In Eq. 11, *r_ik_* denotes the center-to-center distance between the two amino acids. To take into account the polarity of the amino acids, we introduce a binary vector of dimension *N_p_* = 3, such that

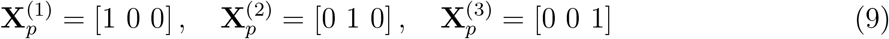

where **X**^(1)^ represents a non-polar amino acid, **X**^(2)^ represents an uncharged polar amino acid, and **X**^(3)^ represents a charged polar amino acid. In addition, we also added another component to the net vector, which is a real value representing the percentage of the buried part of the amino acids (%SASA_buried_), which is defined as the ratio of the buried surface with the solvent accessible surface area of the amino acid in the protein structure, and it is represented by the vector **X**_4_. Note that vector **X**_4_ can also include other properties, such as the temperature, concentration of the salt and pH value of the experiment; therefore, we can write:

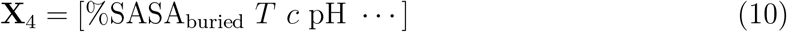

where *T, c*, and pH are the temperature (in kelvin), concentration (in molar) and pH, respectively.

To determine the descriptor vector of the macromolecule, such as protein, we concatenate the vectors **X**_1_, **X**_2_, 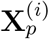 and **X**_4_ into a net descriptor vector **X** with length *N* = 55. Note that in the expression given by Eq. 11, other properties can be encoded. For example, we can encode the dielectric properties in the vicinity of each amino acid in the structure by modifying Eq. 11 as follows:

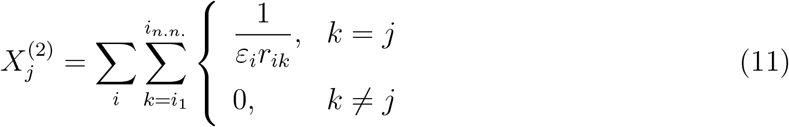

where *ε_ik_* is the dielectric constant of the environment in the vicinity of the mutated amino acid *i*, which can be taken a simple distant dependent dielectric constant between the amino acid *i* and its nearest neighbor *k*: *ε_ik_* = *Dr_ik_*, where *D* is a constant, or even other complicated distance dependence functions.^32,33^ However, in this work, other more complicated distance dependent dielectric constant is considered, such as the sigmoidal function:^32,33^

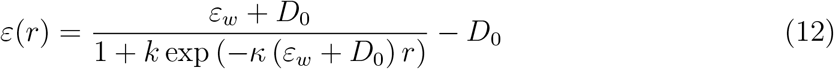

where *r* is the distance between two amino acids, *ε_w_* = 80 is the dielectric constant of water, *D*_0_ = 8, *κ* = 0.5/(*ε_w_* + *D*_0_), *k* = (*ε_w_* − *ε_p_*) / (*D*_0_ + *ε_p_*) with *ε_p_* = 2 being the dielectric constant of protein. A plot of the *ε*(*r*) versus the distance r is presented in Figure 3 for both simple function of the distance of dielectric constant and sigmoidal distance dependence function of the dielectric constant. Here, sigmoidal function gives a smooth variation of the dielectric constant screening the electrostatic interactions from 2 (which is the dielectric constant of the internal protein) to 80 (which is the dielectric constant of bulk water, as shown in Figure 3.

**Figure 3:**
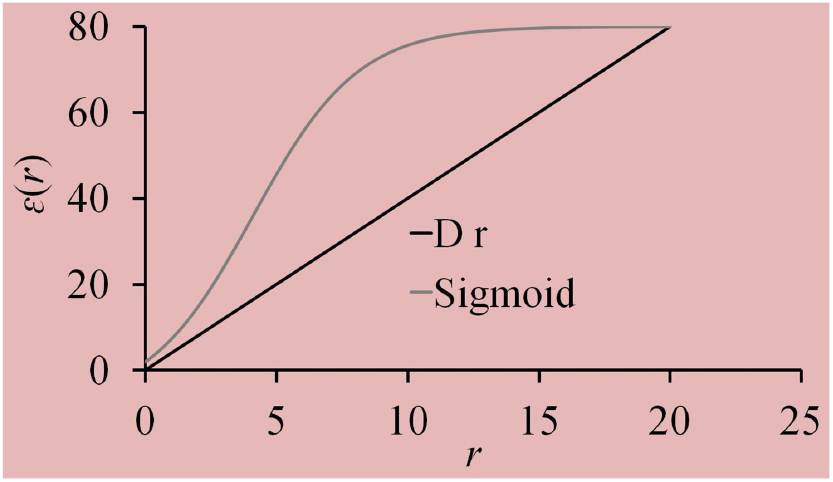
Dielectric constants as a function of the distance r between the amino acids for the two cases: *ε* = *Dr* with *D* = 8, and dielectric constant given by Eq. 12 for *ε_w_* = 80 is the dielectric constant of water, *D*_0_ = 8, *κ* = 0.5/(*ε_w_* + *D*_0_), *k* = (*ε_w_* − *ε_p_*) / (*D*_0_ + *ε_p_*) with *ε_p_* = 2 being the dielectric constant of protein.

Note that these fingerprints of the structures are rotation and translation invariant. Furthermore, as a sequence of the amino acids in a macromolecular structure, the protein data bank, RCSB PDB,^34^ can be used that is unique. Therefore, the descriptor vector **X** is unique representation of a macromolecule in a dataset. In addition, the descriptor vector **X** has the same length for any set of the macromolecules used as input.

It is important to note that if the chemical sample space of the input descriptor vector becomes quite large, then the so-called *principal components analysis* ^35^ can be performed to reduce the degrees of freedom.

## 3 Web-based Services

### 3.1 Structure

Figure 4 shows a flowchart of the offered web-based services. It consists of several databases using different experimental or quantum mechanics data. In particular, it includes the hydration free energies of molecules database;^12^ the heat of formation (or standard enthalpy of formation) of small molecules using quantum mechanics calculations with the Perdew-Burke-Ernzerhof hybrid functional (PBE0).;^5,20^ pKa values experimentally calculated for different amino acids in different proteins;^14–17^ the experimental changes on the Gibbs free energies of the mutant proteins. ^18,19^

**Figure 4:**
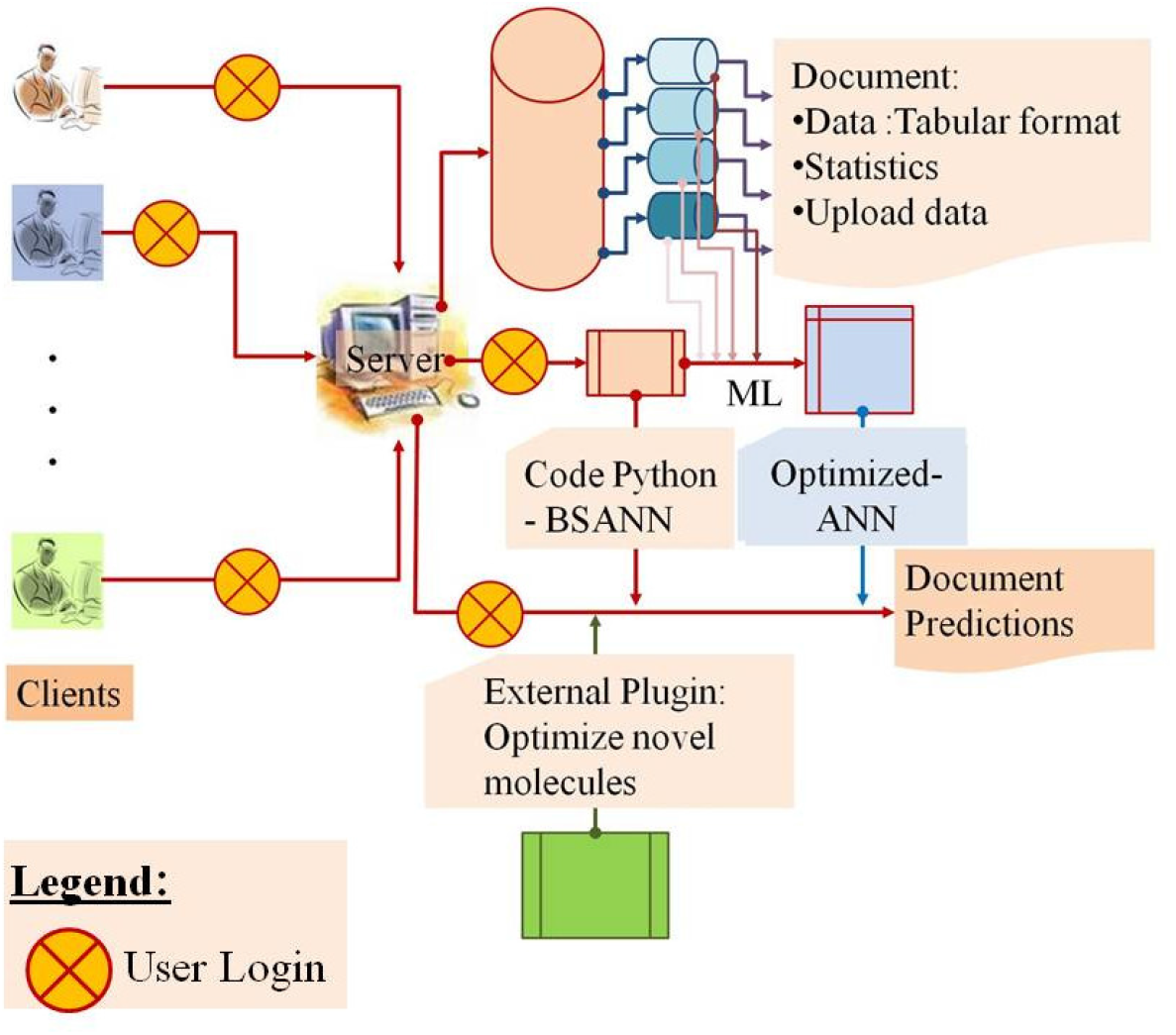
The flowchart of the web based services.

The *clients* are the personal computers (PCs), where the users with a login account will be able to access the first layer of the provided web-based services from the main computer, namely the *server*. With that first session login, the users will be allowed to access and manipulate the data from different databases. In particular, the users can read the data from each database in a tabular form, which then can be copied and pasted in the local computer. Besides, the users can see some statistics about the data included in each dataset, allowing for a judgment of the distribution of the data. Also, the users are allowed using the forms provided on the web to upload new data in a particular database. These data will initially be marked as “not checked”, but after the main administrator of the web-based services verifies the authenticity of the information, the data will be added to the existing database. That can allow the databases to increase the amount of information in the future.

Using the data for each dataset, the main administrator, frequently, performs the optimization of the artificial neural network parameter using an exhaustive machine deep-learning approach by employing the BSANN python code, internally in the server. Note that only specific users are also allowed to perform the optimization on the server using special Login information. Then, the second session of login, for specific users, is allowed to use other services of our tools. In particular, those specific users can use web-based services online to design and optimize novel molecules. Then, using the optimized artificial neural network parameters with the training data, they can predict different properties provided for these molecules. It is interesting to note that the external plugins use graphical interfaces, making the services very user-friendly.

### 3.2 Datasets

There are four databases served using that web-based service. The first database contains 642 small organic molecules, for which we know the experimental hydration free energies,^12^ which has also been subject to our previous studies.^36^ The second database contains 1475 experimental pKa values of inozable groups in 192 proteins both wild type (153 proteins) and mutated (39 proteins).^14–17^ The third database has 2693 experimental values of the Gibbs free energy changes in 14 mutant proteins.^18,19^ The last database has 7101 quantum mechanics heat of formation calculations^5^ (and the references therein), the so-called *QM7*, which is a subset of the so-called *GDB13* molecules, optimized at the quantum mechanics level with the Perdew-Burke-Ernzerhof hybrid functional (PBE0).^20^

In Figure 5, we show the distribution of the average experimental pKa for each residue in the wild-type and mutated proteins of the database.

**Figure 5:**
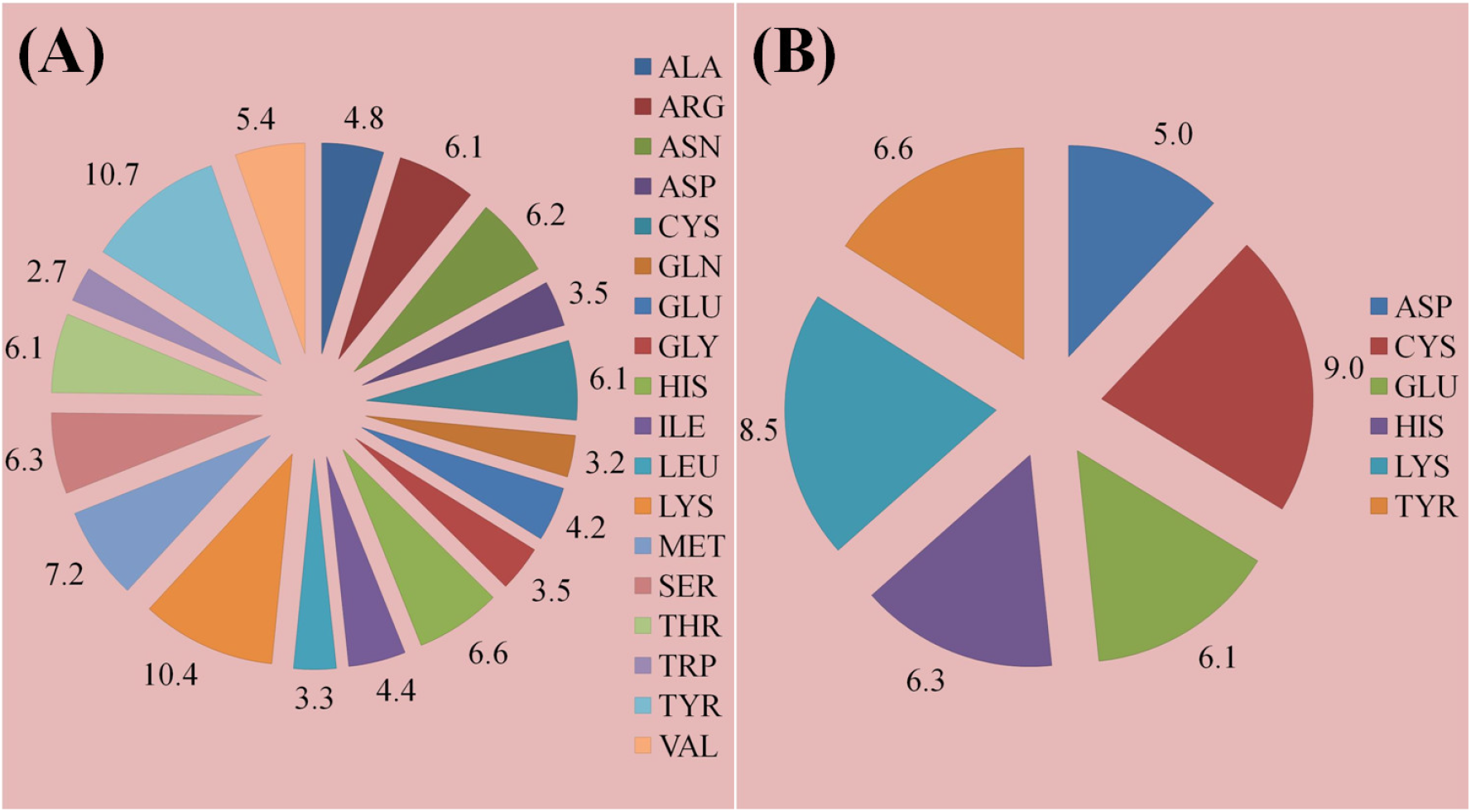
Average experimental pKa for each residue in the wild-type (A) and (B) mutants proteins of the database.

In Figure 6, we show the distribution of the percentage of each type of mutation in the database for which we know either ΔΔ*G* or ΔΔ*G*^H_2_O^, namely 1: single mutation; 2: double mutations, and so on, up to 6: six mutations.

**Figure 6:**
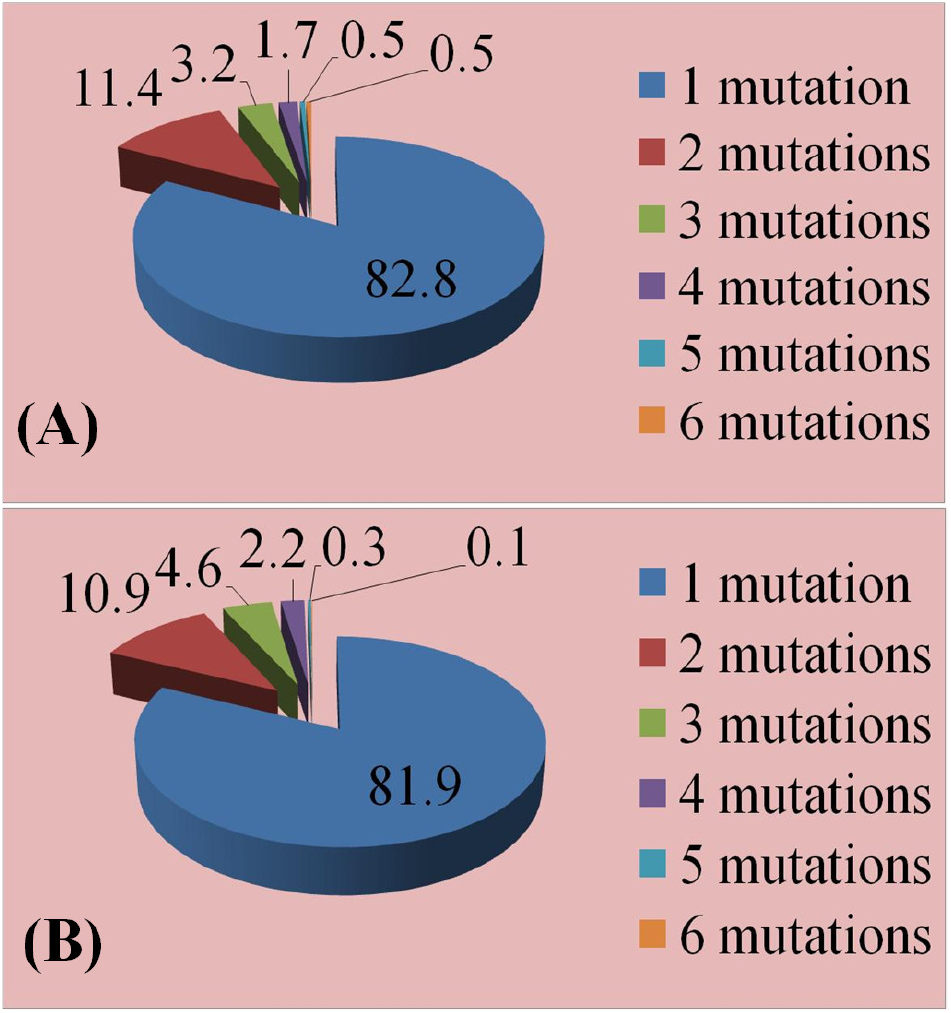
Percentage of each type of mutation in database for which we know either ΔΔ*G* (A) or ΔΔ*G*^H_2_O^ (B). 1: single mutation; 2: double mutations, and so on, 6: six mutations.

All the data are prepared and optimized in advance using AMBER force field^21^ in CHARMM macromolecular computer simulation program. The BSANN code performing the optimization and prediction is in Python computer programming language. The descriptor vectors of the small molecules are based on the Coulomb matrix and the sum over bonds properties, and for the macromolecular systems they take into account the chemical-physical fingerprints of the region in the vicinity of each amino acid.

Software implementing the methods discussed in this study is free for download from the website “https://github.com/kamberaj/bsann”. Also, from the same website, the database of all molecular structure and topology files, used in our calculations prepared using general AMBER force field, can be accessed via the web-based service.

## 4 Results

In this section, we show some results of the predictions using different datasets. For that consider a method to learn a function from a finite dataset 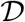 of input-output pairs, namely (**X**, **Y**), where **X** is the feature descriptor input vector for each atom and **Y** is the reference output vector for each atom, such as the hydration free energy, change on the Gibbs free energy, heat of formation, amino acid pKa, and so on. The dataset is then split into a training dataset 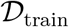 used for learning (or gaining experience) and a validation dataset **D**_valid_ used for testing the knowledge, such that

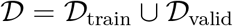

In this study, we will discuss the ability of the training dataset to optimize the parameters of the artificial neural network as a function of the size of the training dataset. In particular, we intend to determine optimal average interval of the confidence such that the error in prediction of a value is small; that is, determining the average confidence interval within which we can predict with a certain confidence level (such as 95 %) the value of any new validation data.

It is important to note that “optimal” here does not the necessary mean small value of the average interval. Instead, an optimal average interval is one that provides the highest value of the confidence level, within which most of the predicated values lie in. In the following discussion, we will use the term “match” for such cases, that is, when the prediction interval of error coincides with that provided by the experimental value using statistical confidence of 95 %, which is verified using the so-called *Student t-Distribution*.

To determine the average confidence interval, we used the following statistical justification. ^37^ Suppose that there are *N*_train_ training data points, and n is the number of the neural networks defining the bootstrapping model given above. Then, the 100(1 – *α*) % bootstrapping confidence interval of the average value for each data point prediction is given by:

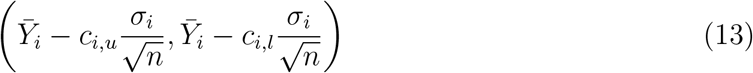

where *σ_i_* is the unbiassed standard deviation obtained from the bootstrapping data distribution. *c_i,u_* and *c_i,l_* are the upper and lower critical values, respectively, determined from the empirical distribution function *F* of the bootstrapping dataset as:

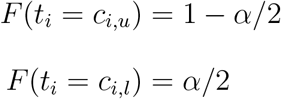

where *t_i_* is the studentized bootstrapping random variable obtained from the data points of the ith prediction.^37^ Then, the average statistical error from all prediction *N*_train_ data points is calculated using the chain rule as follows:

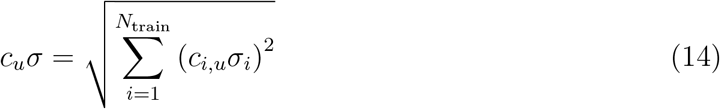

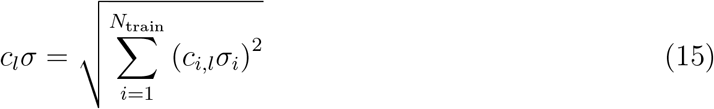

where it is assumed that the statistical errors obtained for each of the training data points are independent, which is indeed the case. Then, the average 100(1 – *α*) % bootstrapping confidence interval of the average value for each data point prediction is defined as:

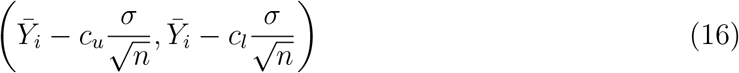

Eq. 16 is used to determine the upper and lower bound of the average values in all our predictions shown in the graphs of the following discussions. Note that in practice, the Student *t*-distribution can be used to approximate the distribution of the bootstrapping dataset, and hence *c_l_* = –*c_u_* = *t*_*n*−1,*α*/2_, which is the critical value for the t-distribution. In this study, the confidence level was *α* = 0.05.

### 4.1 The hydration free energy database

Table 1 summarizes the Pearson coefficient, mean average error (MAE) (in kcal/mol), root mean square error (RMSE) (in kcal/mol) and Matches (in %) for different lengths of training dataset. Predictions are based on the neural network parameters optimized using only the training dataset. Our results show that an MAE value as small as 1.192 kcal/mol is obtained in the validation dataset, corresponding to a Pearson coefficient of 0.886 and an RMSE of 1.770 kcal/mol, for a size of the training dataset of 350 molecules (or equivalently, 84 % of the overall dataset). For that case, the percentage of matches between the predicted and experimental values is 81.5 % with an 95 % statistical confidence.

**Table 1:**
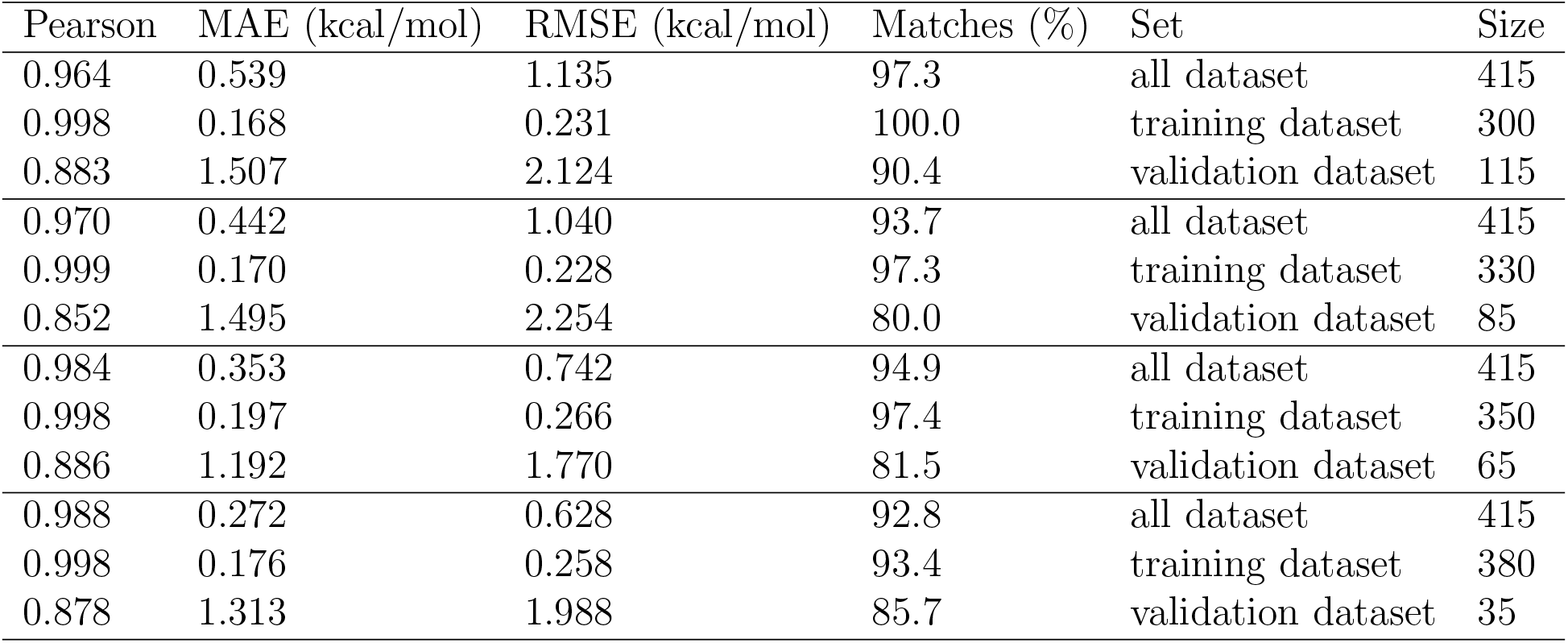
Pearson coefficient, MAE (in kcal/mol), RMSE (in kcal/mol) and Matches (in %) for different numbers of training data. Predictions are based on the neural network parameters optimized using only the training data set.

In Figure 8, we present scatter plots of the experimental hydration free energies and predicted free energies for two different training data sizes, *N* = 330 and *N* = 380 molecules. Errors are calculated using the bootstrapping method and the straight line represents the function *f*(*x*) = *x*. Besides, we have indicated the 95 % bootstrapping confidence interval of error with parallel lines. Results show that a training dataset of size 380 molecules determines better the confidence interval than the training dataset of size 330 molecules. That is because the distribution of the data points of the training dataset influences on the topology of the input data, and thus, larger the input dataset more insights into the structure of the data distribution can be revealed.

**Figure 7:**
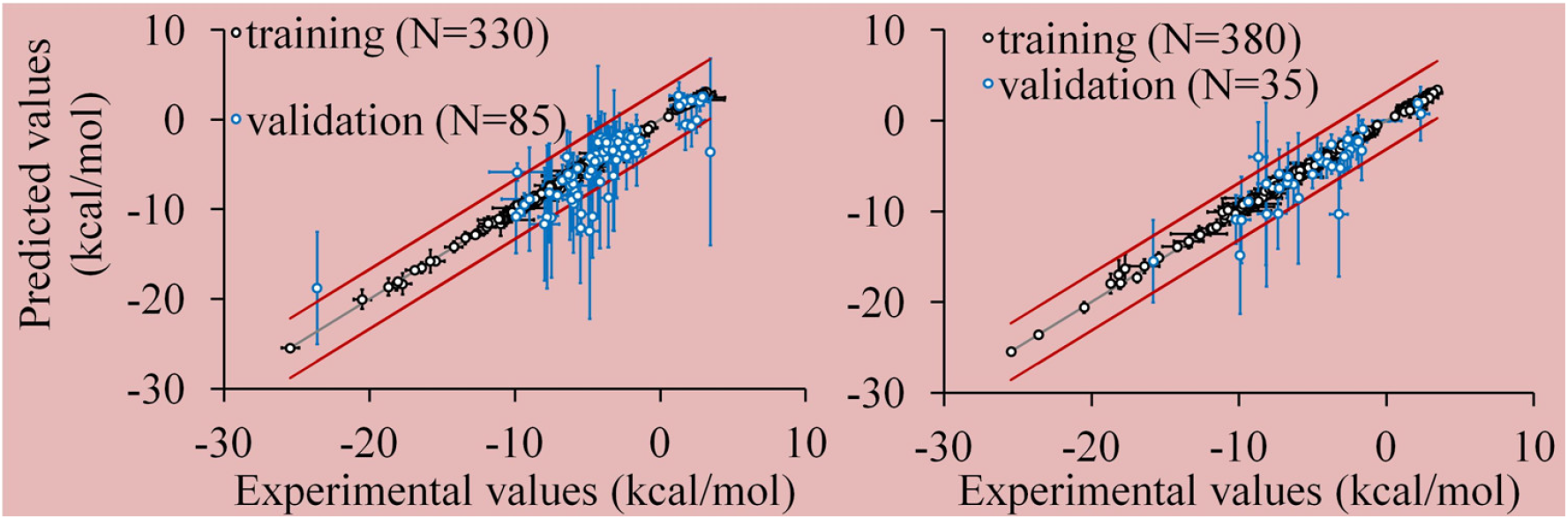
Parity plots between the experimental hydration free energy and predicted free energies. Errors are calculated using the bootstrapping method and the straight line represents the function *f*(*x*) = *x*. The dataset size was 415 molecules. The straight red lines represent the boundary of the 95 % bootstrapping confidence interval of error.

**Figure 8:**
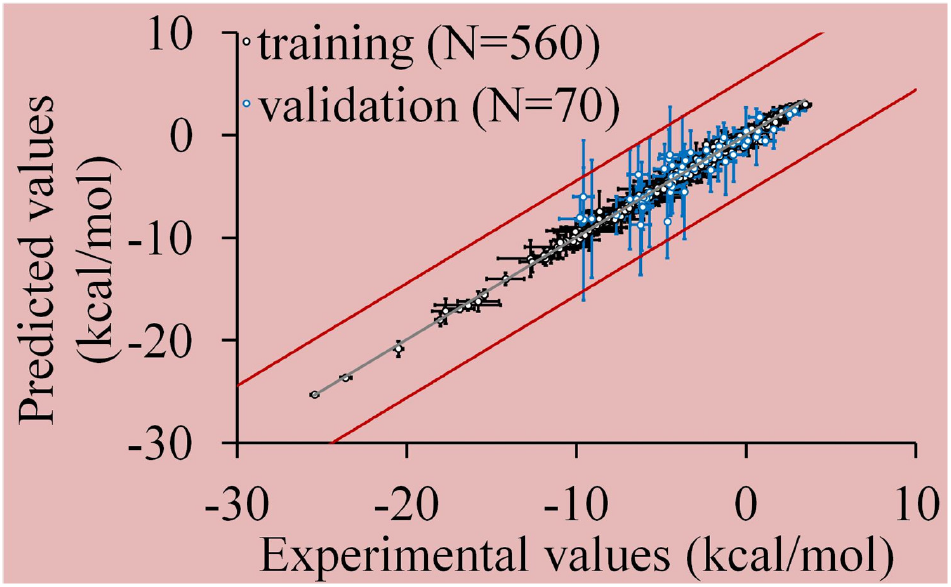
Parity plots between the experimental hydration free energy and predicted free energies. Errors are calculated using the bootstrapping method and the straight line represents the function *f*(*x*) = *x*. The dataset size was 630 molecules. The straight red lines represent the boundary of the 95 % bootstrapping confidence interval of error. For the training dataset: MAE= 0.208 kcal/mol, RMSE= 0.286 kcal/mol and *R* = 0.997; for the validation dataset: MAE= 0.732 kcal/mol, RMSE= 1.050 kcal/mol and *R* = 0.945.

We also used a larger dataset of 630 molecules to train and test the neural network, as shown in Figure 8. For this case, we used 560 (equivalent to 89 % of the total size of the dataset) data points to train the neural network, and the rest about 70 data points for validation (or equivalently, about 11 % of the entire dataset). Our results show an improvement of the predictions when compared with the smaller dataset, as used above; that can be shown by our results presented in Figure 8, indicating that all the validation data points lie inside the 95 % bootstrapping confidence interval. For this dataset, the following values of MAE, RMSE, and Pearson correlation coefficient *R* were obtained: For the training data, MAE= 0.208 kcal/mol, RMSE= 0.286 kcal/mol and *R* = 0.997; for the validation data, MAE= 0.732 kcal/mol, RMSE= 1.050 kcal/mol and *R* = 0.945.

### 4.2 The pKa of amino acids in proteins

Table 2 presents the results of the predictions on both the training and validation datasets. The size of the entire dataset is *N* = 953 pKa calculations. Our results indicate that the Pearson correlation coefficient is above 0.95 in both training and validation dataset. On the training dataset, the smallest MAE and RMSE were 0.104 kcal/mol and 0.164 kcal/mol, respectively. For the validation dataset, the smallest MAE and RMSE values were 0.269 kcal/mol and 0.416 kcal/mol, respectively, obtained for the size of the training dataset about 89 % of the entire dataset. In addition, the matches between the experimental and predicted values of the pKa on the validation dataset is 82 % with a statistical confidence of 95 %.

**Table 2:**
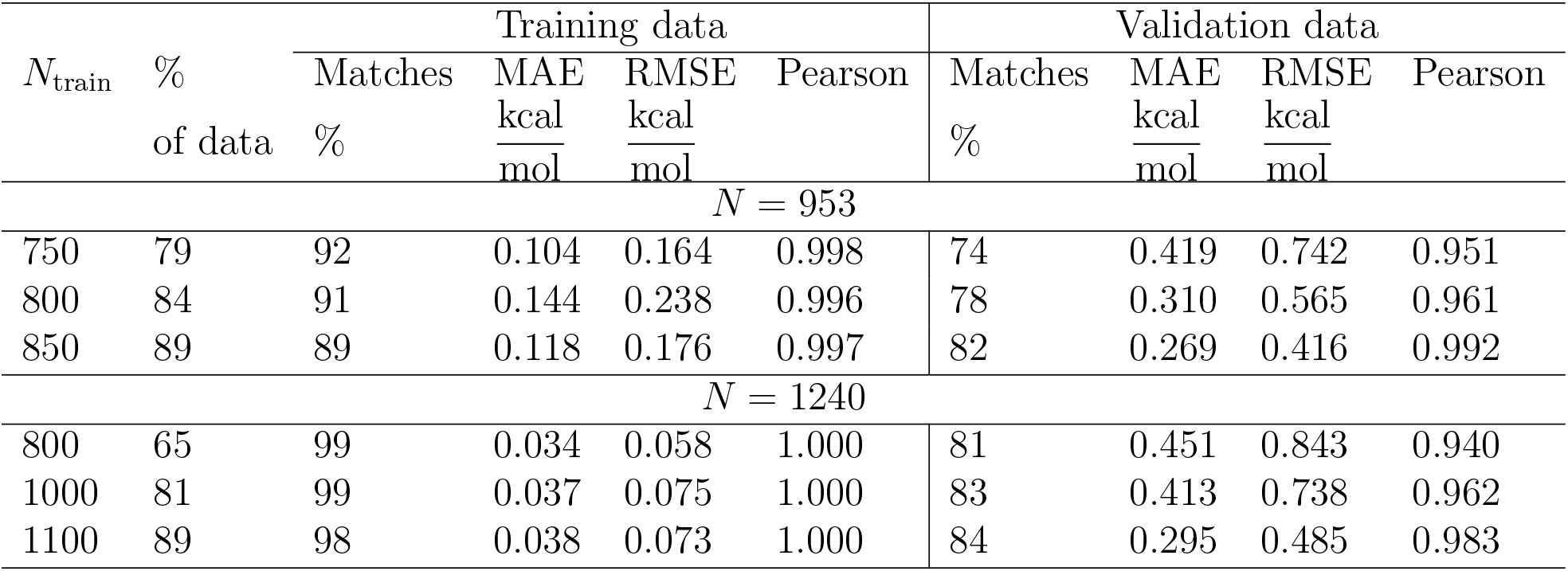
Pearson coefficient, MAE (in kcal/mol), RMSE (in kcal/mol) and Matches (in %) for different numbers of training data. Predictions are based on the neural network parameters optimized using only the training dataset. The sizes of the datasets are *N* = 953 and *N* = 1240 pKa calculations.

In Figure 9, we present the predicated and experimental pKa values graphically as a scatter plot. In addition, we show the average 95 % bootstrapping confidence interval of error of the predicated values within the training dataset. The plots are created for two training data sizes, respectively, 84 % and 89 % of the entire dataset. Interestingly, our results indicate that almost all the validation data predication of pKa values lie inside the average bootstrapping confidence interval of error when 89 % of the dataset is used in training the neural network.

**Figure 9:**
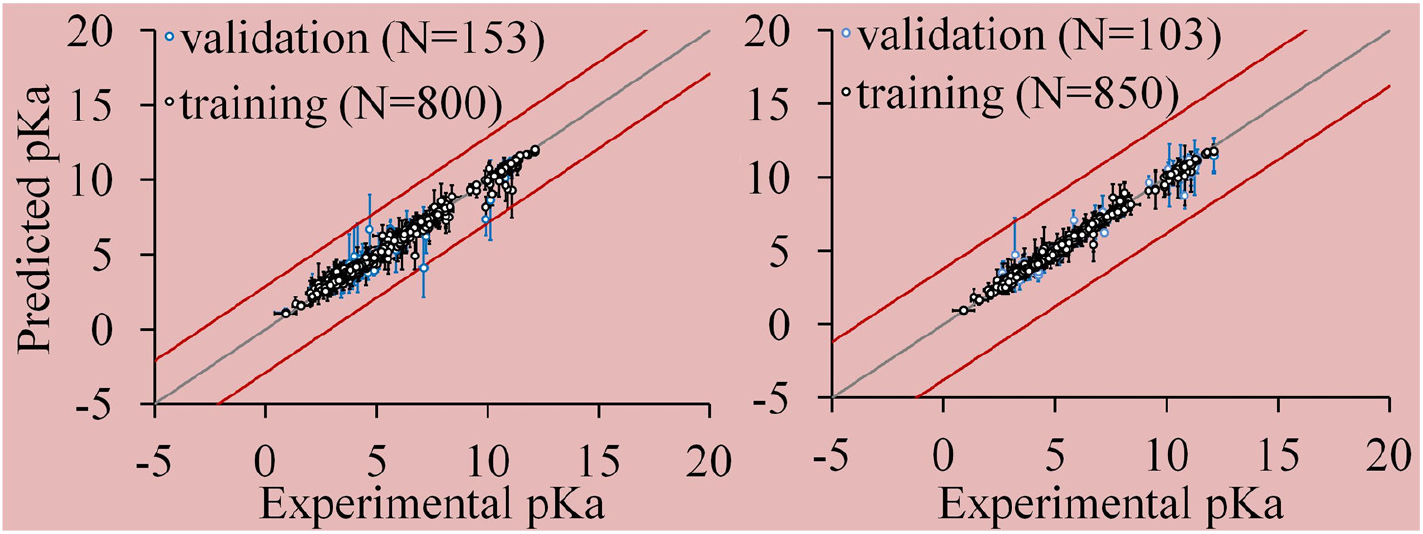
Parity plots between the experimental pKa values and predicted pKa values. Errors are calculated using the bootstrapping method and the straight line represents the function *f*(*x*) = *x*. The size of the dataset was *N* = 953. The straight red lines represent the boundary of the 95 % bootstrapping confidence interval of error.

To check the influence of the dataset size on the learning efficiency from the data, we optimized the neural network for a larger dataset of 1240 pKa calculations. We implemented three different training datasets for the optimizations of the neural network to check the influence of the size of the training data set and the length of the entire dataset, which determine the topology of the input data. We notice that the dataset with *N*_train_ = 1100 (which is about 89 % of the entire dataset) data points for training provided the best optimization. The results are summarized in Table 2, and plotted in Figure 10. Our results show that the MAE decreases about twice for the same percentage of data in the training set taken from a smaller dataset, namely MAE= 0.038 kcal/mol; a smaller RMSE is also obtained for this dataset of about 0.073 kcal/mol. Furthermore, it can be see that the percentage of matches on the training dataset increases to 95 % with a perfect Pearson correlation between the experimental and the predicted of about *R* = 1.000. Moreover, our results show (Figure 9) that the average 95 % bootstrapping confidence interval of error is larger, and all the predicted values of pKa of the validation set lie within bootstrapping confidence interval.

**Figure 10:**
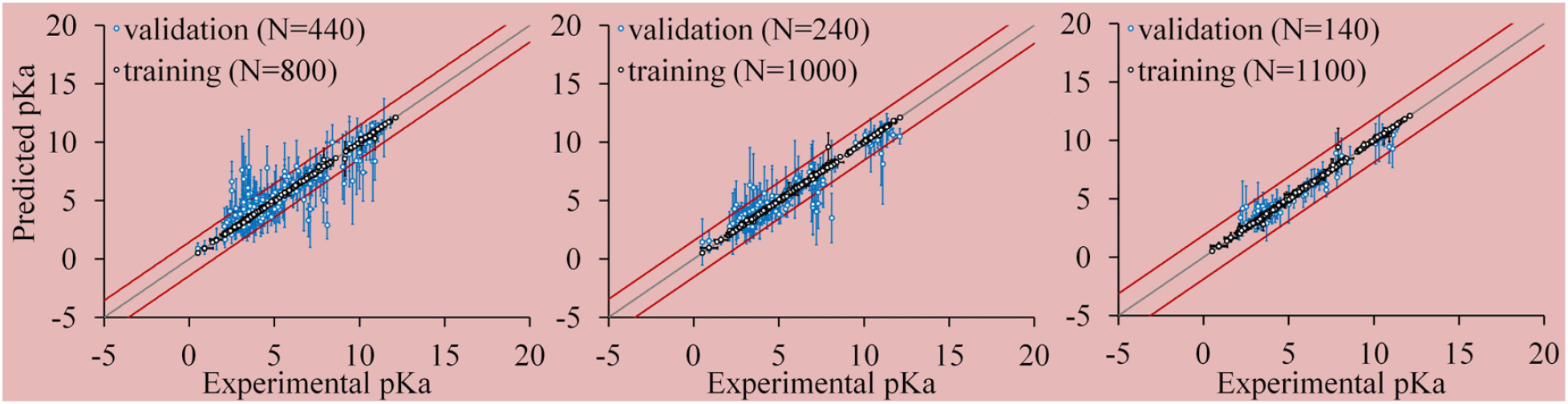
Parity plots between the experimental pKa values and predicted pKa values. Errors are calculated using the bootstrapping method and the straight line represents the function *f*(*x*) = *x*. The size of the dataset was *N* = 1240. The straight red lines represent the boundary of the 95 % bootstrapping confidence interval of error.

### 4.3 Quantum mechanics database

The results of the predicted values of the heat of formation from the quantum mechanics calculations using PBE0 method are shown in Figure 11. The size of the dataset is 7000 molecules. We used two different sets of the training data with lengths 3000 (or equivalently 43 % of the size of the dataset) and 5000 (or equivalently, approximately 71 % of the entire dataset). Our results indicate excellent performance of the predictions using the optimized neural network on the validation data; using just 43 % of the entire dataset for training of the neural network, and the test on the validation data show just a few data are outside the predicted average 95 % bootstrapping confidence interval of error, and when 71 % of the total data are used for training, then all the tested calculations from the validation dataset lie inside the 95 % bootstrapping confidence interval. Interestingly, our results indicate that the optimization of the neural network and hence the increase on the experience for a deep-learning are strongly influenced by the length of the data in the dataset. That explains that the topology of the input dataset plays an important role on the experience gained from the training of the neural network. In the following discussion, we argue that this should be related to the topology of the input data.

**Figure 11:**
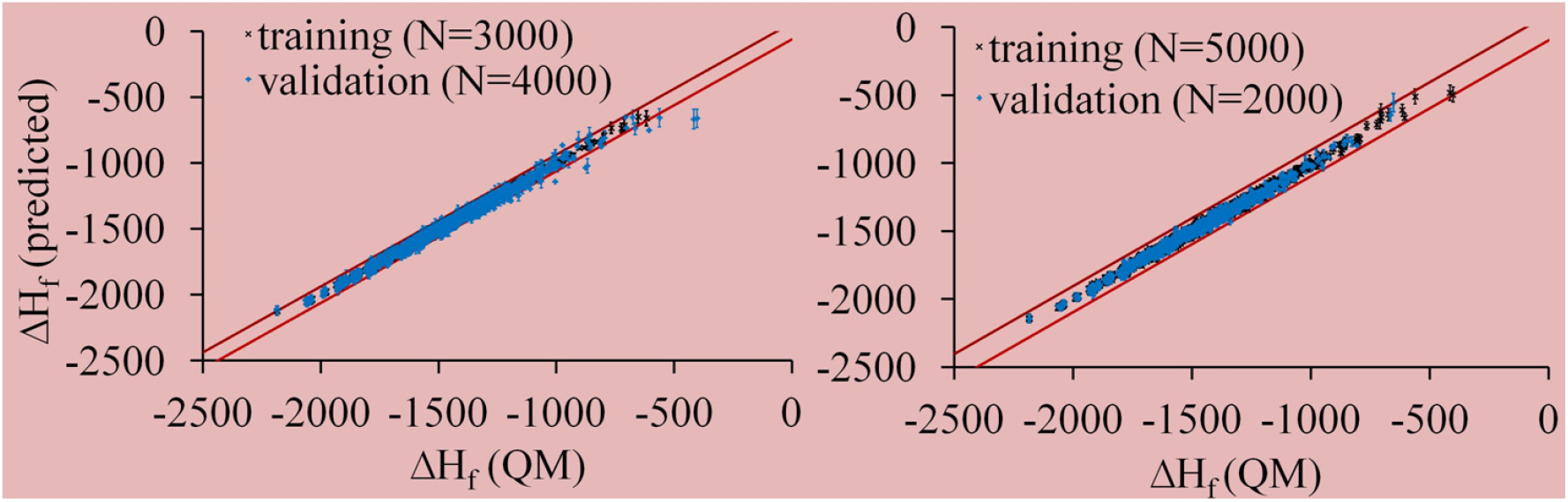
Parity plots between the quantum mechanics heat of formation values and predicted values. Errors are calculated using the bootstrapping method and the straight lines represent the boundary of the 95 % bootstrapping confidence interval of error. Quantum mechanics heat of formation is calculated using the PBE0 method.

## 5 Discussion

In this work, we intend to establish a methodology for an automated machine-like deep learning approach for predicting different (macro)molecular properties. In particular, for a training dataset of molecules, 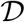 created of *N*_train_ pairs (*X_i_, Y_i_*) for *i* = 1, 2, ⋯, *N*_train_, where the vector **X** denote the feature descriptor vector of dimension *N*_features_ × *N*_train_ and **Y** of dimensions *N*_properties_ × *N*_train_ the reference values. That aims to obtain an estimate of the probability 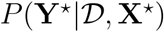 to predict the output 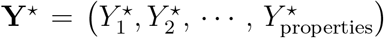 of an optimized neural network for any input test data-point 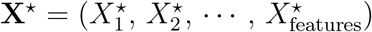. This calculation is now an automated process since the black box is trained to predict the output value described by the probability 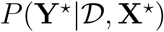, which makes the parameterization of the force fields a very efficient automation process.

However, the accuracy in estimation of 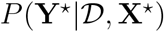 is a data-driven process, and the prediction of **Y**\* depends on the used training dataset. In particular, it depends on the diversity of the feature descriptors for the dataset of molecules, that is, the amount of *N*_features_. Besides, it depends on the size of the dataset, *N*_train_. Both the diversity of the feature descriptors of the compounds and the size of the dataset are interconnected; however, a large size dataset is practically difficult to be established due to the lack of the experimental data, and quantum mechanics data may be expensive to obtain. Besides, increasing the dimensions of the input feature descriptor vector, *N*_features_ × *N*_train_, is equivalent to increasing the amount of information processed by the computer, and hence it increases the amount of the irreversible heat generated during the processing. ^38^ That is related to another computer term, namely “big-data” processing. In the following discussion, we will try to quantify the *weight* of the big-data information by using physical interpretation of information and the principle of the equivalence mass-energy-information. ^39,40^ That will allow us to establish an equivalence between the (necessary) amount of the input training data for accurate prediction of 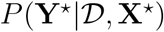 and the limit of the amount of the information that can be processed by a computer considering the heat generated during the computer processing, and so the amount of the external work necessary to process that big-data of information by a computer.

### *Mass* of information

For that, consider that the input feature descriptor vectors are represented in binary form, that is, as zeros and ones. Then, the size of the input matrix of features for all training dataset of compounds is:

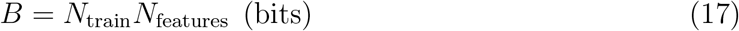

Based on the Landauer principle, ^38,39^ process of adding information into the storage device requires some work done by an external agent:

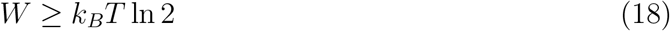

where *k_B_* is the Boltzmann’s constant (*k_B_* = 1.38064 × 10^−23^ J/K) and *T* is the temperature of the environment in kelvin. That work is necessary to change the physical conditions of the environment to add one bit of information into the storage device, which equals the amount of heat generated for creating one but of information in the storage device, and hence:

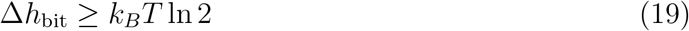

Note that has already been measured experimentally (see, for example, Ref.^40^ and the references therein). Then, the minimum total amount of the heat generated for storing B bits in a storage device at *T* = 300 *K* is

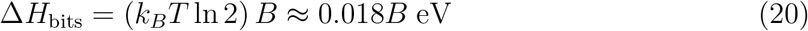

knowing that 1 eV = 1.602 × 10^−19^ J.

For example, suppose that we have 10^4^ training data points and *N*_features_ = 100 for each compound (see also Figure 12), then the total minimum amount of heat generated is:

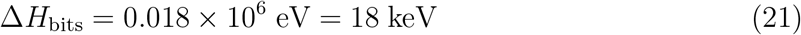

**Figure 12:**
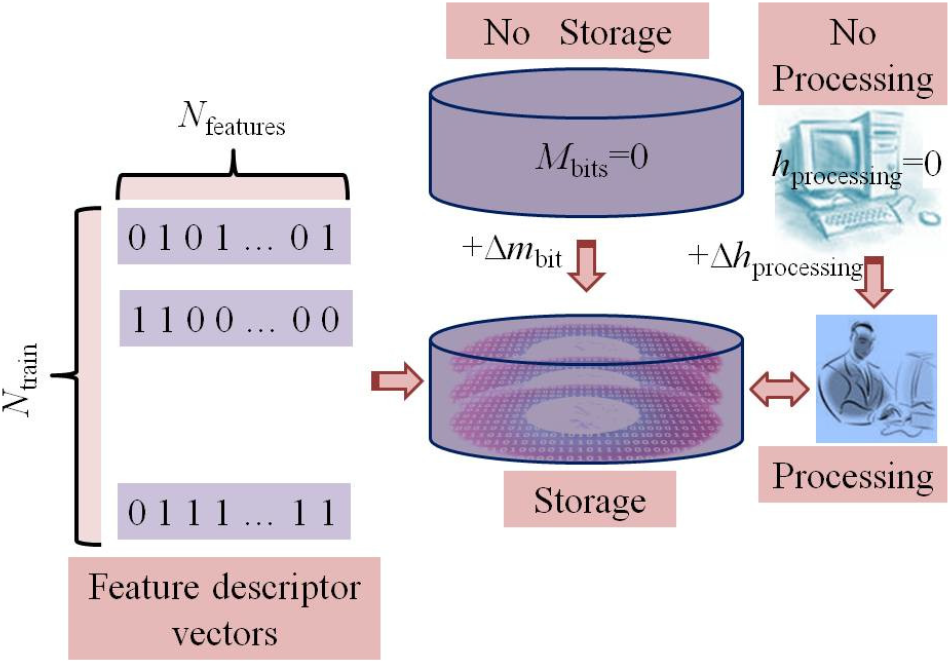
A diagram of the storage and computer processing of the input information. *M*_bits_ gives the mass in kilogram of the amount of information stored in device in bits based on the mass-energy-information equivalence principle;^40^ Δ*m*_bit_ denotes the increase of the mass by adding one bit to the storage device; Δ*h*_processing_ gives the amount of the (irreversible) heat released during the computer processing of the information.^39^

Furthermore, the minimum amount of heat generated for adding one feature (or equivalently, one single bit of information) to each compound of the training dataset is

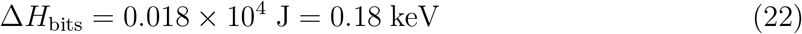

Using the Einstein principle of the mass-energy equivalence, the rest energy of a particle with mass *m* is given

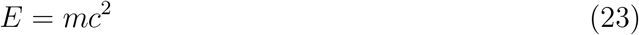

where *c* denotes the speed of light in vacuum, *c* = 3.00 × 10^8^ m/s. Eq. 23 indicates a well-known fact that particles (such as photons) with zero rest mass have zero rest energy. In analogy with mass-energy principle of special theory of relativity, another principles has been postulated of the mass-energy-information equivalence: ^40^

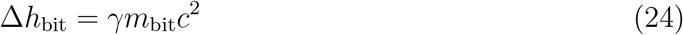

where *m*_bit_ denotes the mass of one bit of information in SI units of kilogram, and *γ* is

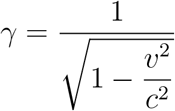

where *v* would represent the speed of storing the information.

Therefore, we have

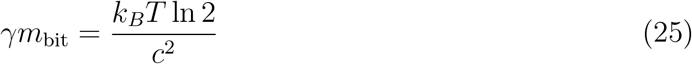

That indicates that when *T* = 0 *K* (absolute zero temperature), then *m*_bit_ = 0 kg. We can assume that absolute zero temperature corresponds to the case when there is no information stored in the device; therefore, for an empty digital storage device *m*_bit_ = 0, and then every added bit of information increases the mass by^40^

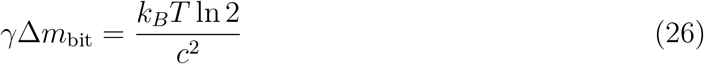

For example, at room temperature of the environment, *T* = 300 K, we have

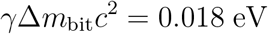

or

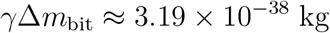

If we consider only the energy of the bit stored in the digital device at rest, then *γ* = 1, and

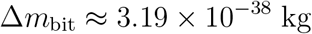

That could give an indication of how fast the bit information moves, the speed of the storing the information in the digital storage device. If we consider again the case of the 10^4^ × 10^2^ = 10^6^ bits of the training data, then the total increase on the mass of the digital device is:

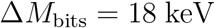

Besides, the increase of the mass of information for adding one bit of information to each input feature descriptor vector of the 10^4^ training data is

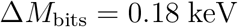

It is interesting to emphasize how beneficial could be if instead of cooling systems to invent a method of how to reuse all that energy released in the form of the irreversible heat.

In addition, one should also consider the energy generated in the form of the irreversible heat by computer processing of that data, as shown in Figure 12. Similar analysis can be followed in that case.

Note that we can also take advantages of new advanced computer architectures to develop new more efficient algorithms. In particular, quantum architecture of the quantum computing can be useful in developing the so-called *quantum artificial neural networks*.^41–43^ In analogy with quantum computing, the quantum bit, the so-called the *qubit*, can be introduced:

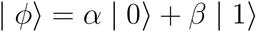

where | 0〉 and | 1〉 are the basis vectors, given as

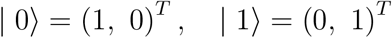

and *α* and *β* are complex numbers such that | *α* |^2^ and | *β* |^2^ give the probability of measuring the state |0〉 and |1〉, respectively, and hence | *α* |^2^ + | *β* |^2^= 1. Based on the quantum artificial neural network model, every input vector can be represented by a qubit, namely | *X_i_*〉, for *i* =1, 2, ⋯, *N*_train_. The output of some layer *j* is then

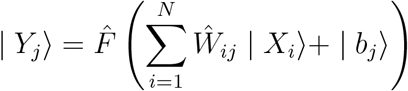

where *N* is the number of neurons, 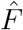 is the quantum operator activation function, 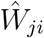 is a 2^*n*^ × 2^*n*^ matrix operator with *n* being the number of qubits acting on the input | *X*_i_〉 (where *n* = 1 in that case, as suggested elsewhere^44^). Here, | *b_j_*〉 qubit denotes the bias term. A learning rule has also been proposed to update the matrix elements 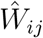 to the desired output qubit | *o_j_*〉 as follows:

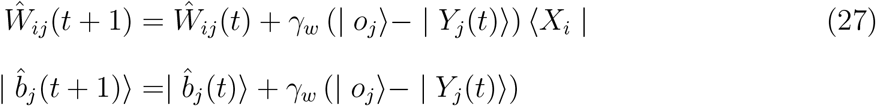

where *γ_w_* is the usual learning constant rate, a number between 0 and 1, and *t* is the iteration step. A comparison of different proposed quantum artificial neural network models is presented in Ref.^44^

Based on the above formalism, then every qubit representing the feature descriptor vectors will increase the mass of the information for 10^4^ training data by

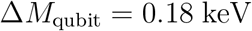

which is 100 times smaller than the classical artificial neural network.

### Topological Data Analysis

It has been argued elsewhere^24^ that the diversity of the feature descriptors of the compound database is essential to increase the range of the test data that can be predicted since the machine learning methodology works very well in interpolating the new data points, but suffers on extrapolating new data outside the range covered by the training dataset. Therefore, one of the critical future developments of the automated machine learning methodologies is the choice of the training dataset and the feature descriptors of the chemical compounds. In Ref.,^24^ a topological data analysis tools is discussed to analyze the feature descriptors of the molecules, which will be introduced in the following. The topological data analysis (TDA) is a field dealing with the topology of the data to understand and analyze large and complex datasets.^45,46^ Here, we are analyzing the dataset represented by a vector of feature descriptors of length *N* and each data point has a dimension *D*:

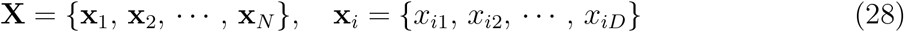

For example, *N* may represent the number of molecules in the dataset and D number of specific features for each molecule. Moreover, as in Ref.,^24^ we assume that the data of the dataset are hidden in a “black box”, for example, a *database*, and also, they are about to be used by a machine learning, which is another “black box”. In such a situation, knowing about the topology of the data *(e.g*., the sparsity of the data points) is of great interest. Note that the TDA is applicable even when the user has access to the data, that is, the structure of the molecules of the dataset is known a priory. In such a situation, the TDA can be applied to determine the topology of the key feature descriptors for each molecule. Note that the TDA is employed to reveal the intrinsic persistent features of the DNA and RNA.^47,48^ Therefore, the construction of the topological spaces upon the input data of a machine learning approach can be applied for each dimension separately, namely to the time series of the form ***X**_d_* = {*x*_1*d*_, *x*_2*d*_, ⋯, *x_Nd_*}, or for each molecular structure, namely ***X**_k_* = {*x*_*k*1_, *x*_*k*2_, ⋯, *x_kD_*}. But, it can also apply to both dimensions at the same time, for instance, by constructing the input data in the form of the following time series obtained by aligning feature descriptors of the molecular structures in one dimension:

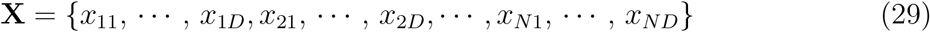

In that case, the input vector of the feature descriptors is a time series of length *N*_train_ = *ND*.

Then, to determine the topological space for this dataset, we first define a distance *σ* > 0. The Vietoris-Rips simplicial complex *R*(***X***, *σ*) or simply Rips complex for each *k* = 1, 2, ⋯ as a *k*-simplex of vertices 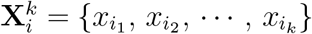 such that they satisfy the condition that the mutual distances between any pair of the vertices is less than *σ*:

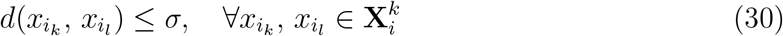

With other words, a *k*-simplex is part of a *R*(**X**, *σ*) for every set of *k* data points that are distinct from each-other at a resolution *σ* and hence the Rips complexes form a filtration of the data from the dataset at a resolution *σ*. That is, for any two values of the resolution *σ′* and *σ* such that *σ* < *σ′*, then

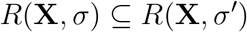

where ⊆ denotes the subset.

All the vertices of a *k*-simplex can be connected in a two-dimensional space by undirected edges forming a graph, which can have different two-dimensional shapes. Figure 13 illustrates how to build simplicial complexes using a set of point cloud data by increasing the resolution value *σ*.

**Figure 13:**
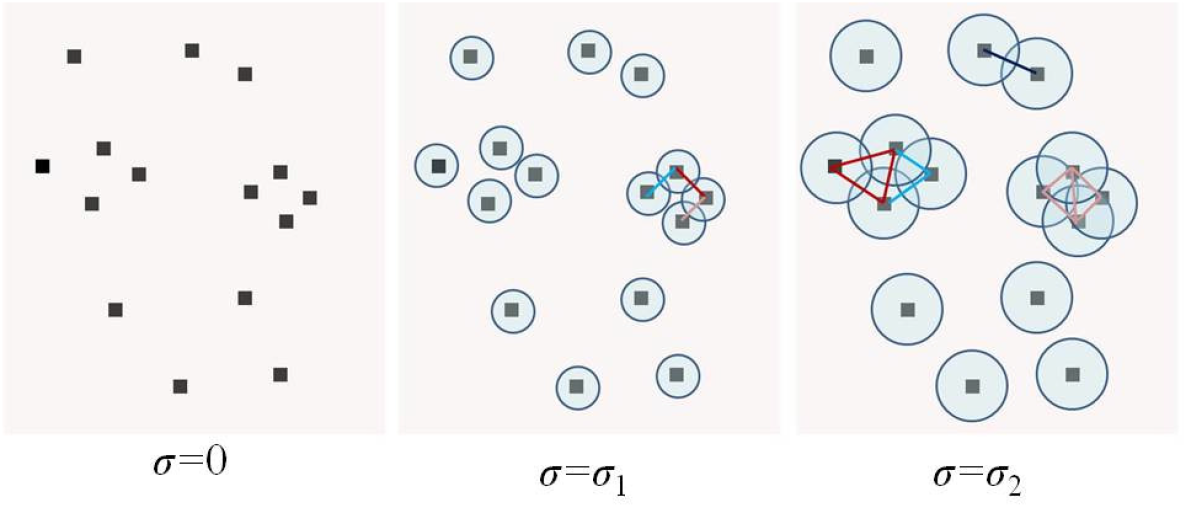
An illustration of building the simplicial complexes by increasing the resolution value *σ*. ^24^

The *k*-simplex dataset points form a loop that is called *hole*. By increasing the resolution *σ*, the shapes grow, and some of the holes die, and some new holes are born. This process is the so-called *σ* loop expansion. The interval between birth and death of a hole is called *persistence interval* indicating whether a hole is structurally relevant or just a noise into the data.

Persistent homology (PH) is an essential tool of TDA, which aims to construct a topological space gradually upon the input dataset, which is done by growing shapes based on the input data. Persistent homology measures in this way the persistence interval of the topological space. The features will be identified as persistent if after the last iteration they are still present.

This procedure is analog to systematic coarse-graining and is of crucial importance for any attempt at capturing natural feature descriptors in terms of a few relevant degrees of freedom, and thus they form the essential philosophical basis of a dataset for the machine learning approaches.

We can argue that the fundamentals of the PH notions on the relevance or irrelevance of perturbations in the data analysis are crucial, and the persistence homology can be considered as necessary as the renormalization group theory in statistical physics when applied to equilibrium phenomena in understanding the relevant or irrelevant interactions. In this analogy, the resolution scaling σ on the topological data analysis can be considered similar to the characteristic correlation length scale that determines the judgment of the strong interactions and correlations renormalization group theory. ^49^

### Feature descriptor vectors in higher dimensions

Note that in our discussion above, the feature descriptor vectors for each compound are considered time invariant, and thus they represent only two- and three-dimensional feature descriptors of the (macro)molecules. However, higher feature descriptor vectors can also be constructed, such as four-dimensional feature descriptor vectors by including the time as the fourth dimension. In that case, to construct the three-dimension part of the feature descriptor vectors, different conformations of the compounds can be taken into considerations, for example, as generated from the molecular dynamics simulations. In that case, the threedimensional configurations of the compounds generated from the simulations can be mapped in a three-dimensional grid, where the centers of the grid points will represent the average positions of each atom obtained from its fluctuations after the configurations are aligned to remove the overall translation and rotation motion of the compounds. Therefore, the feature descriptor vectors obtained from these average structures mapped in a three-dimension grid are translation and rotation invariant. A review of such higher dimensional descriptor vectors is discussed in Ref. ^50^

## 6 Conclusions

In this study, we presented a web-based service for automation of (macro)molecular properties predictions using a new algorithm integrated into a machine learning approach. That web-service is made up of four different databases of both molecular and macromolecular systems properties. Each database has a user-friendly interface that provides the possibility to upload information into the database, which then is verified by the main administrator of the service. Besides, the clients can perform statistics on the web related to each database, obtaining in this way the information contained in each database in a tabular or graph format.

Besides, our web-based service provides other tools and plugins for prediction of the properties of the new (macro)molecular systems using a newly developed deep-learning approach based on the bootstrapping swarm artificial neural network. Furthermore, we showed, in this study, how to create input descriptor vector for the artificial neural network for both small molecules and macromolecular systems. The descriptor features included both the two-dimensional (macro)molecular fingerprints and the three-dimensional structure of the systems. Moreover, we presented a statistical approach of how to estimate the bootstrapping confidence interval of the error.

The application of that new algorithm on our data indicated that the topological spaces of molecular properties description vector on the relevance or irrelevance of perturbations in the data analysis are crucial. Furthermore, we envision that the persistence homology can be considered as necessary as the renormalization group theory in statistical physics when applied to equilibrium phenomena in understanding the relevant or irrelevant interactions. In this analogy, the resolution scaling factor on the topological data analysis can be considered similar to the characteristic correlation length scale that determines the judgment of the strong interactions and correlations renormalization group theory.

## ASSOCIATED CONTENT

### Supporting Information Available

The Web-based Services along with the databases are published on the following website.

## Acknowledgement

The author (H.K.) thanks the International Balkan University for the support.

